# Hypercholesterolemia aggravates in-stent restenosis in rabbits: a mitigating effect of stent surface modification with CD47-derived peptide

**DOI:** 10.1101/2023.02.27.530304

**Authors:** Ilia Fishbein, Vaishali V Inamdar, Ivan S Alferiev, George Bratinov, Menekhem M. Zviman, Anna Yekhilevsky, Chandrasekaran Nagaswami, Kristin L. Gardiner, Robert J Levy, Stanley J Stachelek

## Abstract

**Background:** Hypercholesterolemia (HC) has previously been shown to augment restenotic response in several animal models and humans. However, the mechanistic aspects of in-stent restenosis (ISR) on a hypercholesterolemic background, including potential augmentation of systemic and local inflammation precipitated by HC are not completely understood. CD47 is a transmembrane protein known to abort crucial inflammatory pathways. Our present studies have examined the interrelation between HC, inflammation, and ISR and investigated the therapeutic potential of stents coated with a CD47-derived peptide (pepCD47) in the hypercholesterolemic rabbit model.

**Methods and Results:** PepCD47 was immobilized on metal foil coupons and stents using polybisphosphonate coordination chemistry and pyridyldithio/thiol conjugation. The relative abundance of the surface-associated cells on bare metal (BM) and pepCD47 foils exposed to whole rabbit blood showed a 40% inhibition of cell attachment on pepCD47-modified surfaces. Likewise, cytokine expression analyzed in buffy coat-derived cells cultured over the BM and pepCD47-derivatized foils demonstrated a M2/M1 increase with pepCD47 coating. Hypercholesterolemic and normocholesterolemic rabbit cohorts underwent bilateral implantation of BM and pepCD47 stents in the iliac location. Hypercholesterolemia increased neointimal growth in comparison with normocholesterolemic animals at 4 weeks post-stenting. These untoward outcomes were mitigated in the arteries of hypercholesterolemic rabbits treated with pepCD47-derivatized stents. Compared to NC animals, inflammatory cytokine immunopositivity and macrophage infiltration of peri-strut areas increased in HC group animals, and was attenuated in the arteries of hypercholesterolemic rabbits treated with pepCD47 stents.

**Conclusions:** Augmented inflammatory responses triggered by HC underlie severe ISR morphology in hypercholesterolemic rabbits. Blockage of initial platelet and leukocyte attachment to stent struts through CD47 functionalization of stents mitigates pro-restenotic effects of HC.

## Introduction

Development of ISR in stented atherosclerotic coronary and peripheral arteries is a common complication of stent angioplasty^1^. Even in the era of advanced drug-eluting stents (DES), ISR complicates up to 10% of the coronary interventions^2^, with the numbers being even higher for patients with certain comorbidities^3^. Given the decisive role of dyslipidemia in the progression of atherosclerosis, the significance of hypercholesterolemia for ISR pathogenesis has been scrutinized for a long time, yet the clinical data are controversial^4–6^. The most plausible link between the elevated total and the low-density lipoprotein-associated cholesterol (LDL-C) and ISR is increased systemic and vascular inflammation associated with hypercholesterolemia^7, 8^. Increased uptake of cholesterol by endothelial cells promotes monocyte attachment and transmigration into the subendothelial space via upregulated expression of cell adhesion molecules^9, 10^. Monocytes differentiate into macrophages and produce multiple cytokines that, by paracrine action, amplify the initial damage to healthy endothelium triggered by hypercholesterolemia^11^. Furthermore, the compensatory anti-inflammatory mechanisms mediated by nitric oxide (NO) get depressed because of reduced NO production by the dysfunctional endothelium and its decreased bioavailability due to NO sequestration by reactive oxygen species^12^. In addition, hypercholesterolemic states induce thrombocythemia and activate platelets to adhere to activated endothelium priming it for neutrophil and monocyte ingress^13, 14^. CD47 is a “marker of self” ubiquitously expressed on the cell surface to protect the host from the attack by its own immune system^15, 16^. We have previously shown that CD47^17, 18^, and CD47-derived peptides^19, 20^ bestow protection from platelet and inflammatory cell attachment to the surface of biomaterials. We also demonstrated that the stainless-steel surfaces of stents functionalized with a 22-aminoacid peptide derived from the extracellular Ig domain of rat CD47 are partially protected from platelet and leukocyte association and reduce ISR after carotid artery implantation in a rat model of stent angioplasty^20^. Since this study used healthy normocholesterolemic rats, the results do not entirely reflect the complex relationship between hypercholesterolemia, vascular and systemic inflammation, and the therapeutic effectiveness of CD47-modified stents. The current study aimed to elaborate on the role of inflammation as a link between hypercholesterolemia and the exacerbated response to vascular injury and to study the anti-inflammatory and anti-restenotic effects of CD47-derivatized stents in a clinically relevant animal model.

## Materials and Methods

### Materials, chemicals, and biologicals

316-grade stainless steel foils and tubes were from Goodfellow (Coraopolis, PA, USA) and Microgroup (Medway, MA, USA), respectively. Open cell design stainless steel (316L) stents were obtained from C2D Medical (Ahmedabad, India). Tris (2-carboxyethyl) phosphine hydrochloride (TCEP^.^HCl) and Vybrant™ CFDA SE Cell Tracer Kit were purchased from Thermo Fisher Scientific (Waltham, MA, USA). 4 % glutaraldehyde and 0.1M Sodium Cacodylate buffer pH 7.4 were obtained from Electron Microscopy Sciences (Hatfield, PA). A peptide derived from the extracellular Ig domain of rabbit CD47 (Ac-Gly-Asn-Tyr-ThrCys-Glu-Val-Thr-Glu-Leu-Ser-Arg-Glu-Gly-Lys-Thr-Val-Ile-Glu-Leu-Lys-LINKER-Cys-OH) was custom synthesized by Bachem (Torrance, CA, USA). A terminal reactive cysteine residue was attached to the pepCD47 core sequence through a dual hydrophilic spacer (AEEAc-AEEAc). Polyallylamine bisphosphonate with latent thiol groups (PABT) and polyethylenimine with installed pyridyldithio groups (PEI-PDT) were synthesized in our lab according to the published procedures^21^. All other chemicals were from Sigma-Aldrich. All cell culture reagents were from Thermo Fisher Scientific (Waltham, MA, USA). A cholesterol fluorimetric assay kit was purchased from Cayman Chemicals (Ann Arbor, MI, USA). ELISA kits for rabbit TNFα and IL1β were from G-biosciences (St. Louis, MO, USA). A lipid peroxidase assay kit was from Eagle Biosciences (Amherst, NH, USA). Elastic tissue Verhoeff-van Gieson staining kit was obtained from Sigma-Aldrich (St-Louis, MO, USA). Primary antibodies to rabbit CD68, TNFα, IL1β, and IL6 were from ProSci (Poway, CA, USA), MyBioSource (San Diego, CAS, USA), Bioss (Woburn, MA, USA), and Abbexa (Cambridge, UK), respectively.

### pepCD47 functionalization of stainless steel

Metal samples were washed in isopropanol and chloroform at 55□C to remove impurities and baked in an air atmosphere at 220□C for 30 min. The specimens were then reacted with 0.5% aqueous solution of PABT at 60□C with shaking for 2 hours, followed by washing in DDW and exposure to 10 mg/ml TCEP^.^HCl in 0.1M acetic buffer at 28□C for 10 min to deprotect thiols. The samples were then washed with degassed DDW and reacted in an argon atmosphere with a degassed 1% aqueous solution of PEI-PDT at 37□C for 40 min and thoroughly washed with DDW. Rabbit CD47-derived peptide (0.5 mg in 1 ml of 50% trichloroacetic acid) bearing free thiols was further diluted with degassed dimethylformamide/PBS mixture and reacted in argon atmosphere with thiol-reactive PDT groups on the surface of metal samples at 37□C with shaking for 30 min. The metal specimens were then washed in sterile PBS and stored at 4□C prior to use.

### Chandler loop experiments

PepCD47 surface-modified or unmodified stainless steel tube inserts (10 mm length, 6.3 mm diameter) were placed in succession with 1 cm intervals into the 40-cm sections of 1/4” PVC tubing (Terumo Cardiovascular Systems, Ann Arbor, MI). Ten ml of NC and HC rabbit blood anticoagulated with sodium citrate (0.6% final concentration) was perfused through the tubing for 3 hours at a shear rate of 20 dyn/cm^2^. At the end of the perfusion period, the steel inserts were removed, gently washed with PBS, and processed, determining cell and platelet adhesion to the metal substrate by CFDA staining and SEM, as detailed below.

#### CFDA staining

To visualize the platelets and inflammatory leukocytes attached to the adluminal surface of the inserts, non-fixed specimens were exposed to 10 μM Vybrant™ CFDA SE at 37°C for 15 mins. The samples were then thoroughly washed with PBS and fixed in 4% paraformaldehyde at room temperature for 1 hour. The tubes were then cut longwise, inverted, and imaged using a fluorescence microscope (Nikon TE300). The samples were also analyzed fluorimetrically (ex 485 nm, em 538 nm, cut-off 530 nm) using Spectra Max Gemini EM (Molecular Devices, Sunnyvale, CA).

#### Scanning electron microscopy

After removing from the Chandler loop apparatus, pepCD47-functionalized and non-modified stainless-steel inserts exposed to HC blood were gently washed with PBS and fixed in 2% glutaraldehyde in sodium cacodylate buffer with 0.1M sodium chloride for 24 hours. The inserts were then cut longwise, inverted, rinsed with DDW, dehydrated serially with 70-100% ethanol and hexamethyldisilazane, and sputter-coated with gold palladium. Ten digital micrographs (magnification range 500-10000) at random areas were acquired for analysis using SEM (Quanta250, FEI, Hillsboro, OR). Platelet and leukocyte attachment in 4 random fields was determined by manual counting.

### Macrophage polarization on the pepCD47-modified and non-modified metal substrate (PCR studies)

Platelet-depleted buffy coats from NC and HC rabbit blood were seeded in the wells of a 12-well plate containing 15 mm × 15 mm stainless steel foil coupons, either non-modified or pepCD47-functionalized (n=4 for each group). The medium was changed after 4 hours for complete RPMI-1640 supplemented with 100 ng/ml of human MCSF that was shown to support the growth and differentiation of rabbit macrophages^22^. The medium was changed every other day. After 6 days of the culture, cells growing on the cell culture plastic (control) and those growing on the different steel substrates were lysed. RNA was isolated using an RNeasy kit (Qiagen, Germantown, MD, USA) and reverse transcribed in cDNA with TaqMan reverse transcription reagents (Applied Biosystems, Waltham, MA, USA). cDNA was amplified in RT-PCR reaction with Sybr Green and rabbit inflammatory cytokine primers (IDT, Coralville, IA, USA), reflecting the M1 and M2 polarization phenotypes.

### *In vivo* studies

All animal experiments were pre-approved by the University of Pennsylvania IACUC and conformed to all relevant regulations. Some animals were fed an HC diet (a standard rabbit chow supplemented with 1% cholesterol and 3% peanut oil) (Bioserv, Flemington, NJ, USA). Blood for monocyte isolation was harvested from the central ear artery. For stent surgery, in isoflurane-anesthetized rabbits of either gender (3-4 kg) vascular access was surgically obtained with arteriotomy in the right carotid artery. Under fluoroscopic control, a 4-Fr sheath was advanced over the 0.035” guidewire to the distal aorta. The guidewire was exchanged for a 0.014” guide wire, over which a 3-Fr Fogarty catheter was inserted in the left common iliac artery. A balloon of the Fogarty catheter was inflated and passed across the artery three times to denude the endothelium, after which the Fogarty catheter was exchanged for a 3.0 mm angioplasty catheter loaded with a stent. The stent was deployed in the segment of the iliac artery devoid of the side branches at 12-14 atm to achieve a stent/artery diameter ratio of 1.3. The guidewire was then relocated into the right common iliac artery, and the sequence of Fogarty balloon denudation/stent deployment was repeated. In the NC rabbits, both stents were BMS, while in the HC rabbit group, one of the stents was BMS, and the other was a pepCD47 stent. All animals were euthanized 28 days after stent placement.

### Statistics

Data are presented as means ± SD, unless specified otherwise. Differences between the groups were analyzed by ANOVA followed by a post-hoc Tukey’s test. Statistical was assigned at p < 0.05.

## Results

To study the attachment of bloodborne cells to the native and pepCD47-modified metal substrate, the relative number of cells adhered to the stainless-steel samples was quantified using CFDA staining and fluorimetry after 3-hour exposure to NC and HC rabbit blood recirculated in a Chandler loop apparatus. While no significant differences between the non-modified and pepCD47-modified samples was noted with NC blood (Fig. 1 A, B, and E), the number of cells on the metal surface exposed to HC blood was reduced by 35% as a result of CD47 functionalization (Fig. 1 C, D and E). These results were further confirmed by scanning electron microscopy that demonstrated a 3.2-fold decrease in the number of platelets from HC blood deposited on CD47-modified steel surface (Fig. 1 F-H). The presence of pepCD47 fostered M2 polarization of the macrophages derived from the buffy coats that originated from NC and HC rabbit blood as evidenced by increased M2/M1 cytokine gene expression ratio in macrophages cultured on the bare metal and CD47-functionalized surfaces (Fig 2). The expression of signature M1 cytokines, IL1β and IL12, was suppressed in macrophages growing on the CD47-modified surface compared to macrophages cultured on bare metal steel (Fig. 2 A and B). In contrast, the M2 cytokines, IL10 and TGFβ demonstrated opposite kinetics (Fig. 2 C and D).

**Fig. 1.**
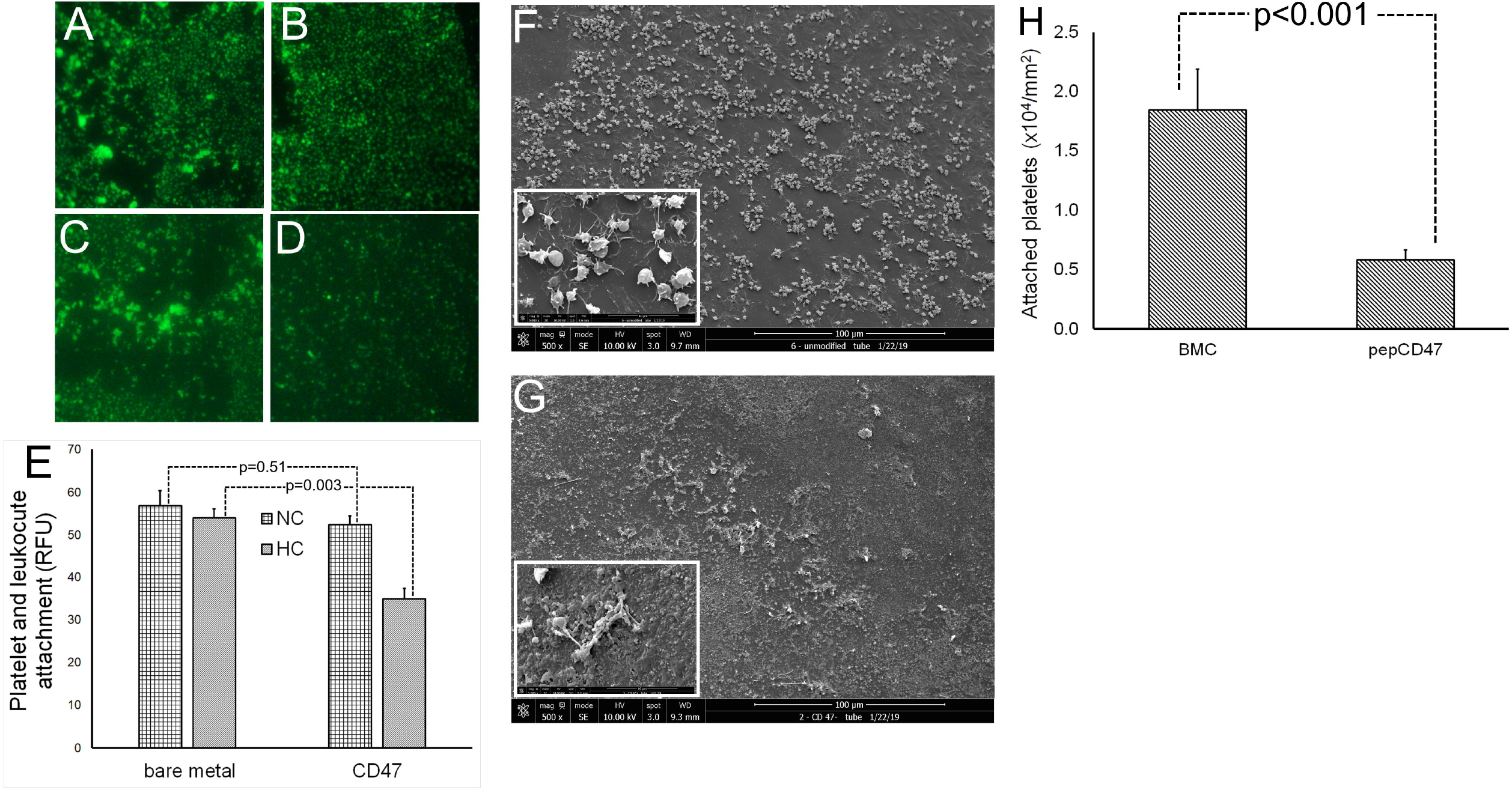
Blood interaction with pepCD47-modified and non-modified stainless steel surface in the Chandler loop flow experiments. (A-D) Fluorescence microscopy images (FITC filter set; original magnification – 200×) of adluminal surface of non-modified (A, B) and pepCD47-modified (C, D) stainless steel inserts exposed to NC (A, C) or HC (B, D) rabbit blood recirculating in the Chandler loop apparatus (n=4 per sample type, blood type). E. Fluorimetry measurement of the relative fluorescence emitted by CFDA SE stained cells associated with the adluminal surface. (F-G) Scanning electron microscopy images (original magnification – 500×; insets – 5,000x) of the adluminal surface of non-modified (F) and pepCD47-modified (G) stainless steel inserts exposed to HC rabbit blood recirculating in the Chandler loop apparatus (n=3 per sample type). (H) The surface density of attached platelets.

**Fig. 2.**
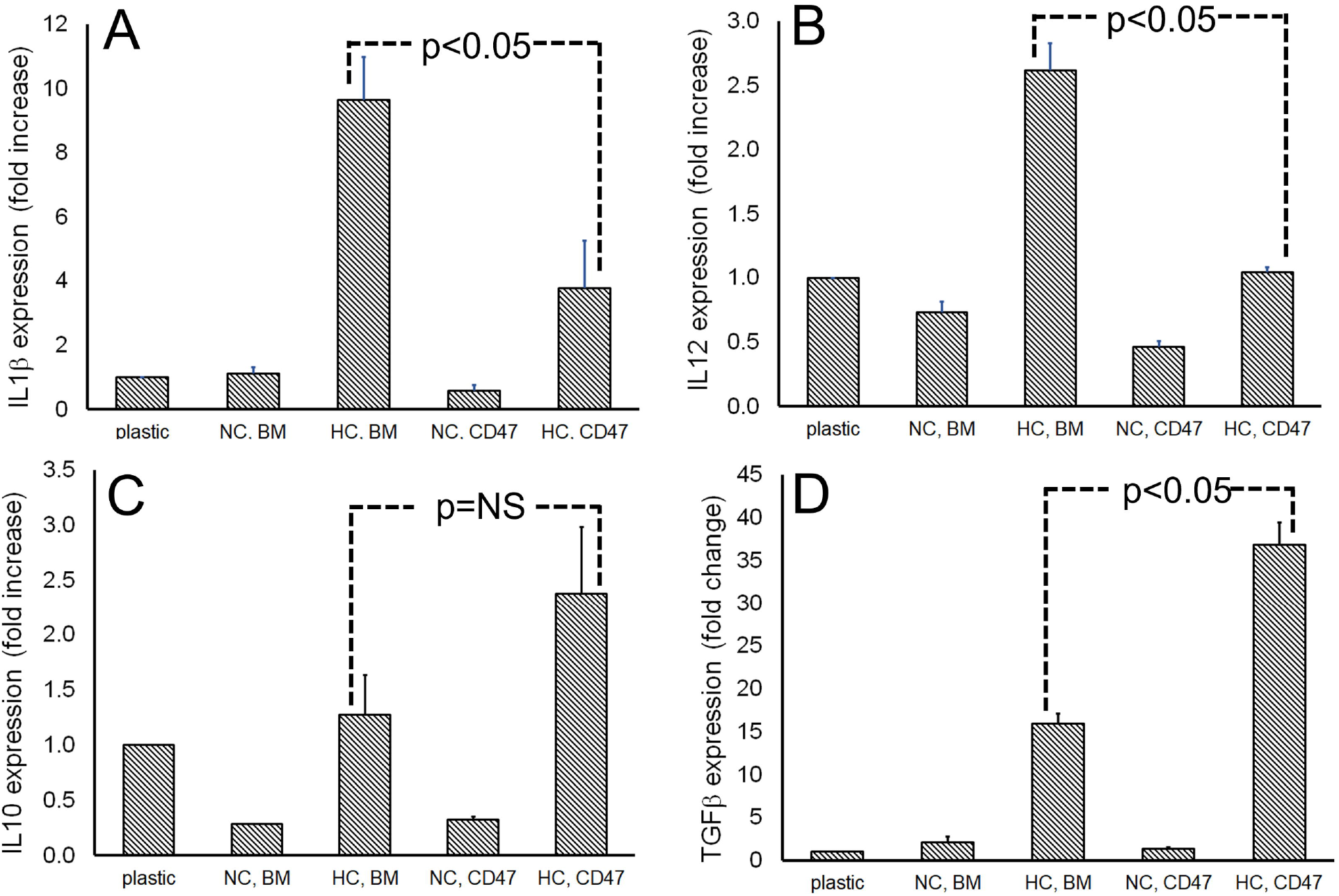
Relative expression of M1 (IL1β and IL12) and M2 (IL10 and TGFβ) macrophage polarization markers by macrophages growing on the bare metal and pepCD47-modified surface. (A-D) Relative quantification (RQ) by 2^-δδct^ method of IL1β (A), IL12 (B), IL10 (C) and TGFβ (D) expression by rabbit NC and HC blood-derived macrophages grown on the cell culture plastic (control), bare metal surface (BMS) and pepCD47-modified steel (n=3 per condition).

To investigate the impact of hypercholesterolemia on ISR development and the antirestenotic effectiveness of CD47 stent functionalization, BMS and stents modified with pepCD47 were implanted in rabbits fed normal chow and HC diet. Exposure to a hypercholesterolemic diet (4 weeks before the intervention and 4 weeks after stent deployment) considerably increased neointimal formation in the stented iliac arteries (Fig. 3 A and B). Neointimal area (Fig. 3 D), neointimal thickness (Fig. 3 E), percent of luminal stenosis (Fig. 3 F), and neointima-to-media ratio (Fig. 3 G) increased 2.3-5.2 fold in the hypercholesterolemic animals. These hyperplastic responses to stent implantation were reversed 29-52% (p<0.01 for all comparisons) in the arteries treated with the pepCD47-derivatized stents (Fig. 3 A-G).

**Fig. 3.**
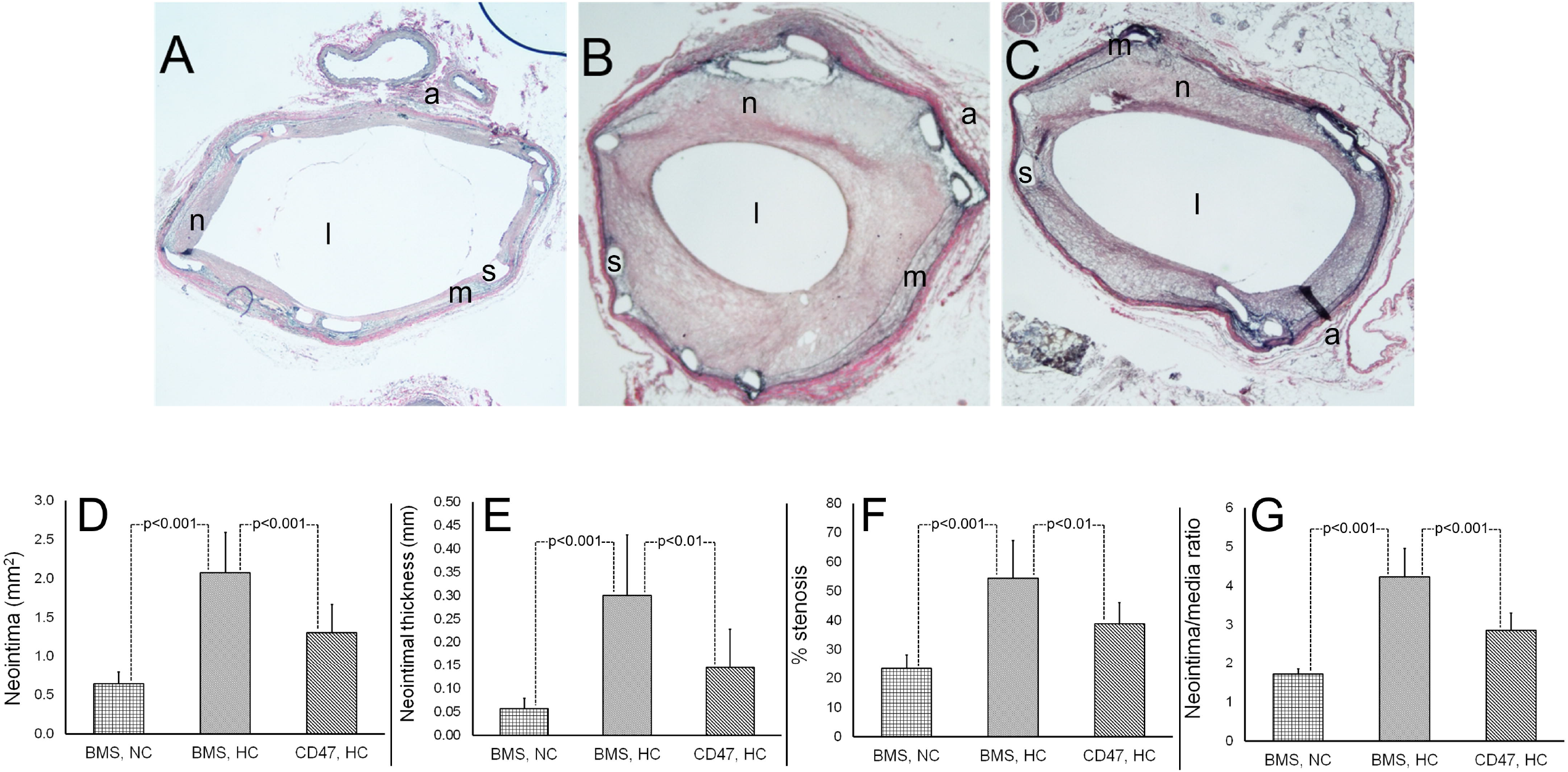
Effects of hypercholesterolemia and stent surface functionalization with pepCD47 on the extent of ISR. (A-C) Representative microscopy images of stented iliac arteries of NC (A) and HC (B, C) rabbits treated with bare metal (A, B) and pepCD47 stents (original magnification is 20x). Stent struts were removed by acid treatment prior to paraffin embedding^35^, sectioning and Verhoef-van Gieson staining. n – neointima, m – media, a – adventitia, l – lumen, s – strut positions. (D-G) Morphometric values of neointimal area (D), neointima thickness above the struts (E), percent of luminal stenosis (F), and neointima to media area ratio in the treatment groups.

In the individual animals treated with BMS, the severity of ISR expressed as neointima to media ratio strongly correlated (Pearson coefficient of 0.826; p<0.0001) with total cholesterol levels in rabbit blood (Fig. 4). BMS implantation in hypercholesterolemic animals was associated with increased plasma concentrations of inflammatory markers, TNFα (Fig. 5 A) and IL1β (Fig. 5 B), and lipid peroxidation marker, malondialdehyde (Fig. 5 C) compared to normocholesterolemic rabbits. Likewise, local arterial expression of inflammatory markers, TNFα (Fig. 6 A, B, and D), IL1β (Fig. 6 E, F, and H), and IL6 (Fig. 6 I, J, and L) was significantly upregulated in the peristrut regions of hypercholesterolemic rabbits compared to normocholesterolemic counterparts. Furthermore, inflammatory macrophage infiltrates were significantly more predominant in the peri-strut regions of hypercholesterolemic rabbits as compared to normocholesterolemic animals (Fig. 6 M, N, and P). Local expression of inflammatory markers and the presence of macrophages around the struts was significantly reduced in the arteries of hypercholesterolemic animals treated with pepCD47-functionalized stents (compare Fig. 6 B and C, F and H, J and K, N and O). Macrophages accumulating in the peri-strut location appear to present the principal source of pro-inflammatory cytokines, as apparent from dual immunofluorescence staining of macrophages and IL1β-producing cells (Fig. 7). Activation of quiescent cells of the vascular wall by the macrophage-released cytokines drives and sustains cell proliferation, migration, and ECM production, thus contributing to the neointimal formation and ISR^23^. In BMS-treated hypercholesterolemic animals, proliferation in the neointimal, medial, and adventitial compartments increased compared to normocholesterolemic counterparts (Fig. 8 A, B, and D) and was significantly reduced in the arteries treated with pepCD47-modified stents (Fig. 8 B, C and D).

**Fig. 4.**
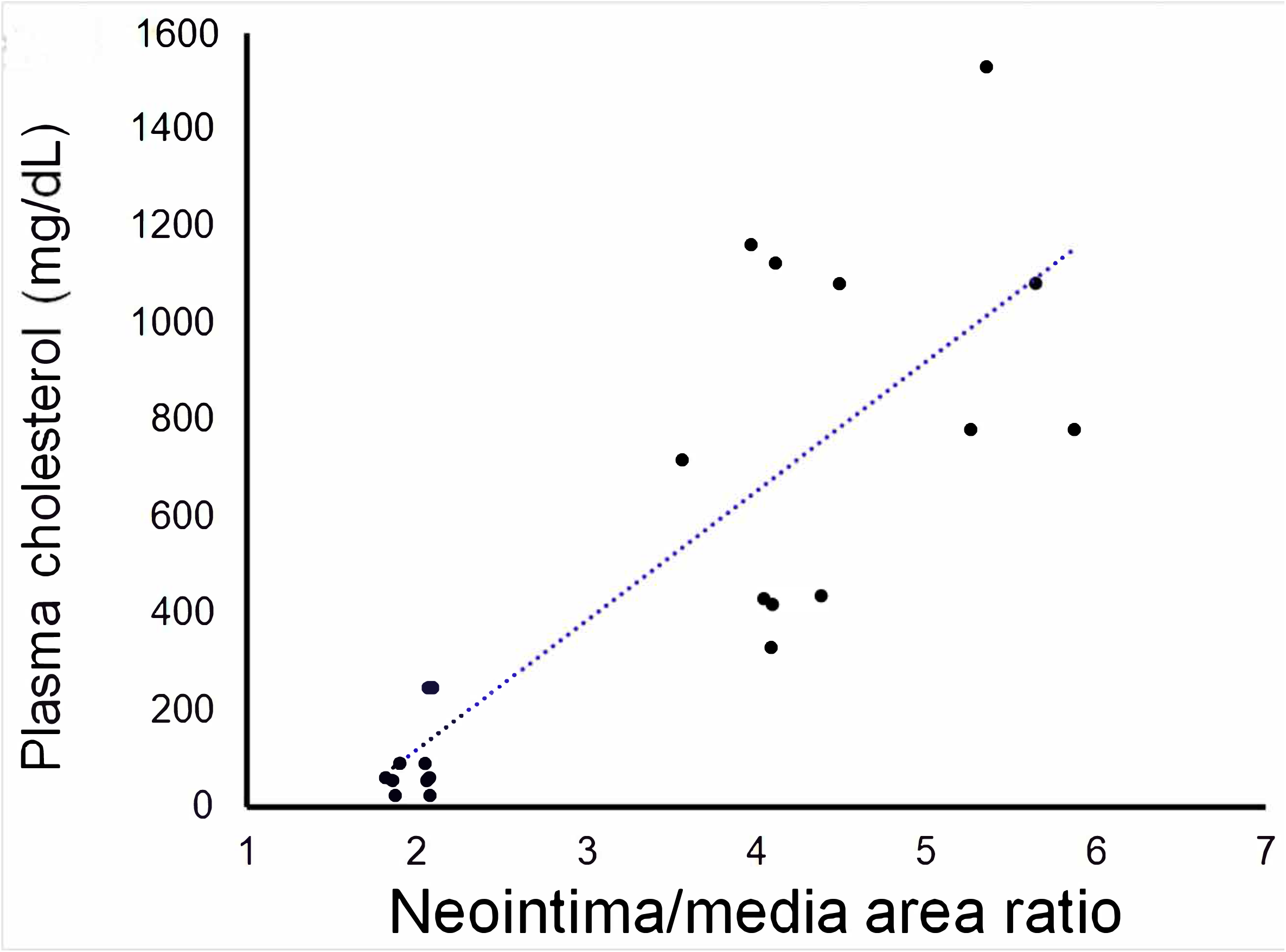
A correlation between the individual plasma cholesterol concentrations and the severity of ISR. Individual levels of plasma cholesterol at euthanasia and the respective N/M ratio values are plotted.

**Fig. 5.**
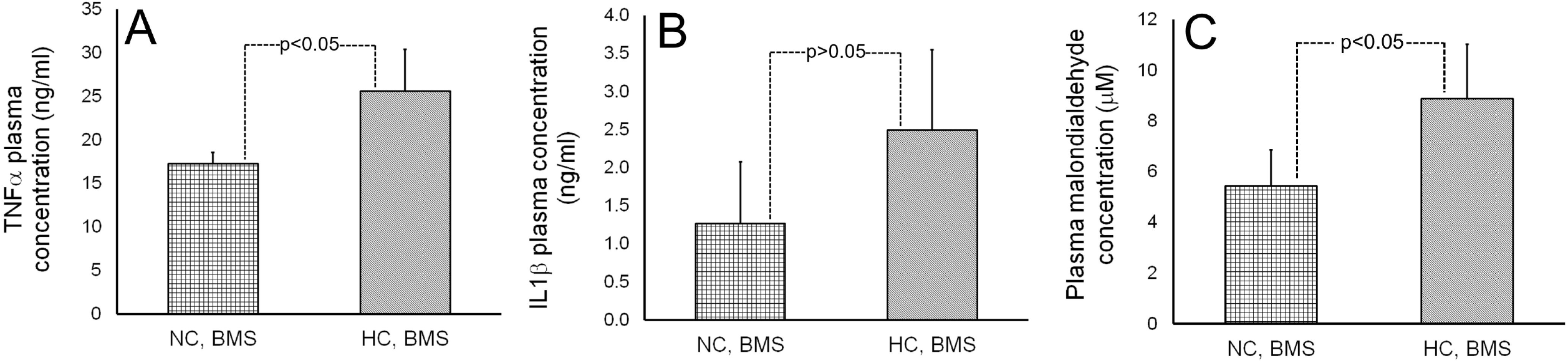
Plasma concentrations of inflammatory and oxidative stress markers in NC and HC rabbits. (A-C) Concentrations of TNFα (A), IL1β (B), and malondialdehyde (C) as determined by ELISA in plasma of NC (n=5) and HC (n=5) rabbits.

**Fig. 6.**
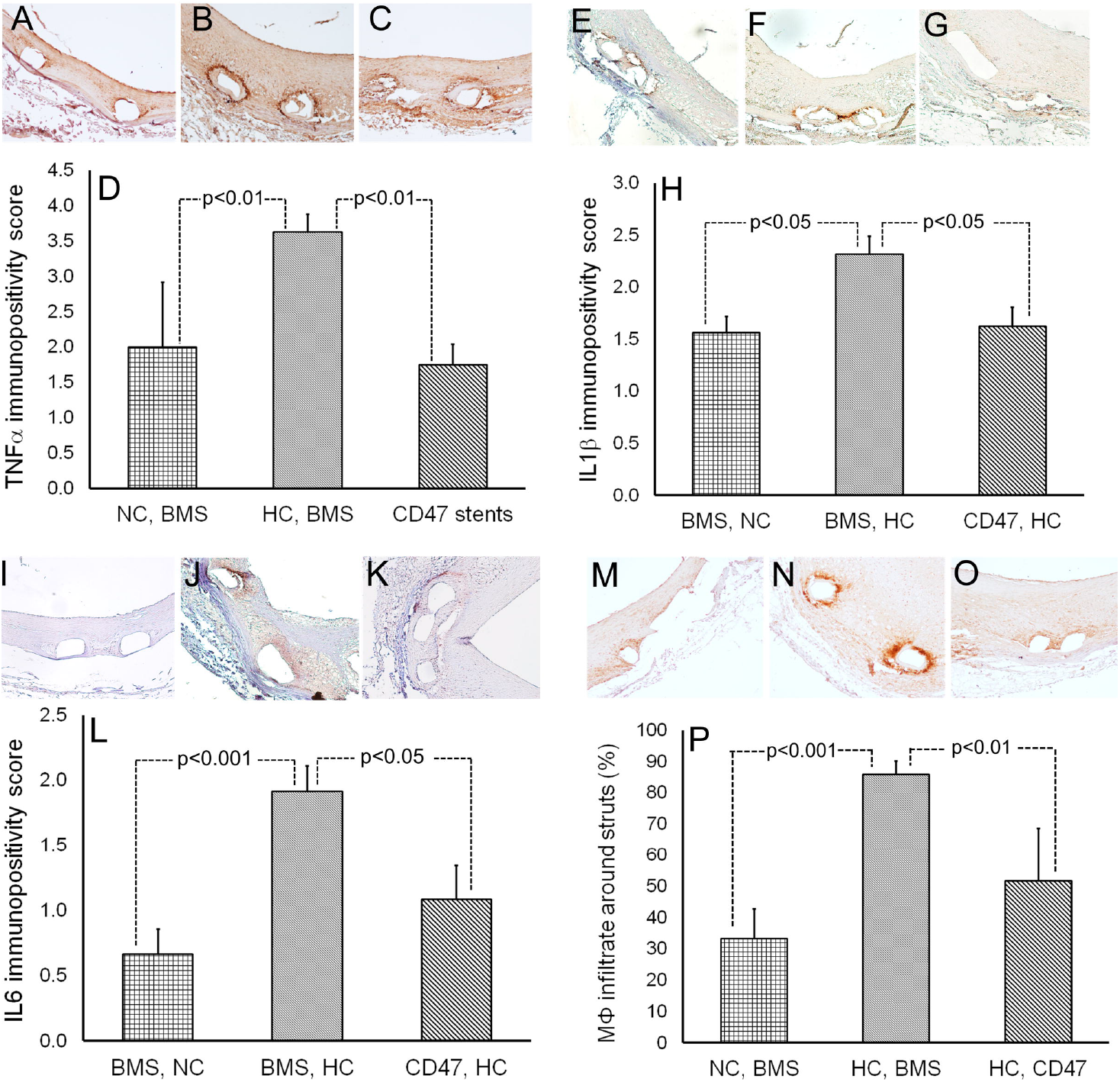
Immunohistochemical assessment of the inflammatory cytokine expression in the peri-strut areas of NC and HC rabbits implanted bare metal and pepCD47-modified stents. (A-P) Representative microscopic images (100x magnification) of iliac arteries stented with BMS (A, B, E, F, I, J, M, N) and pepCD47-functionalized stents (C, G, K, O) and immunostained for TNFα (A-C), IL1β (E-G), IL6 (I-K) and CD68-positive macrophages (M-O). Quantification of TNFα (D), IL1β (H), and IL6 (L) expression was based on a semi-quantitative (0-4) scale. Macrophage infiltration around the struts (P) was calculated as a percentage of a strut circumference occupied with the CD68-positive cells (n=4 for each marker/treatment group combination).

**Fig. 7.**
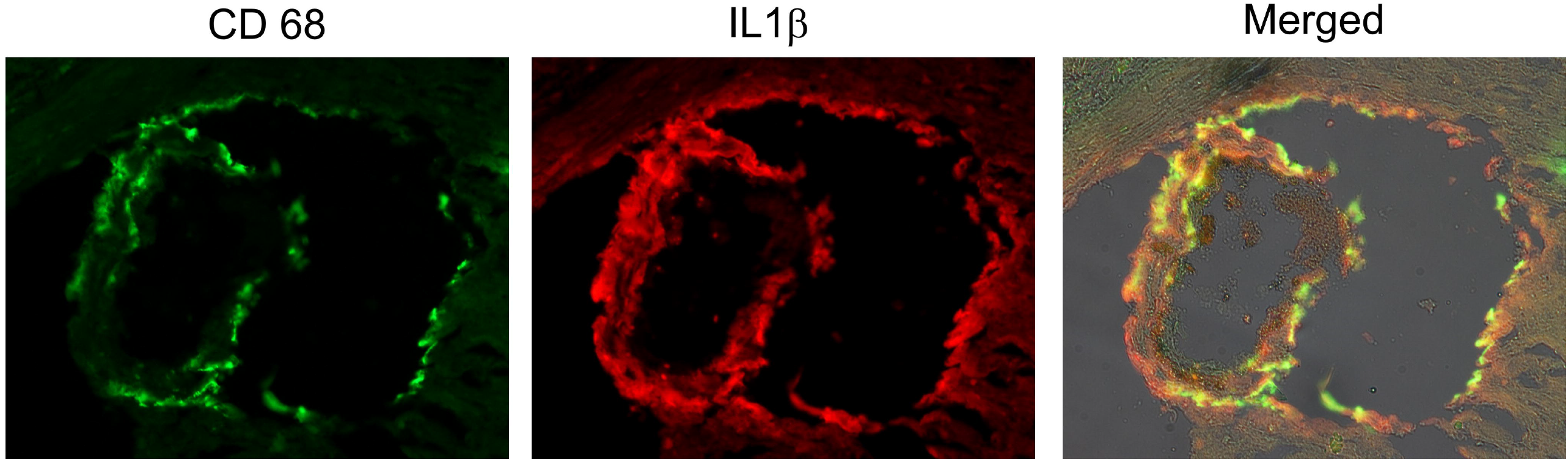
Co-localization of inflammatory cytokine-producing cells with macrophages. (A-C) Microscopic images (original magnification – 200×) of stented arterial section co-stained with anti-CD68 (A) and anti-IL1β (B) antibodies. Merged image (C) demonstrates spatial colocalization of CD68 and IL1β expression.

**Fig. 8.**
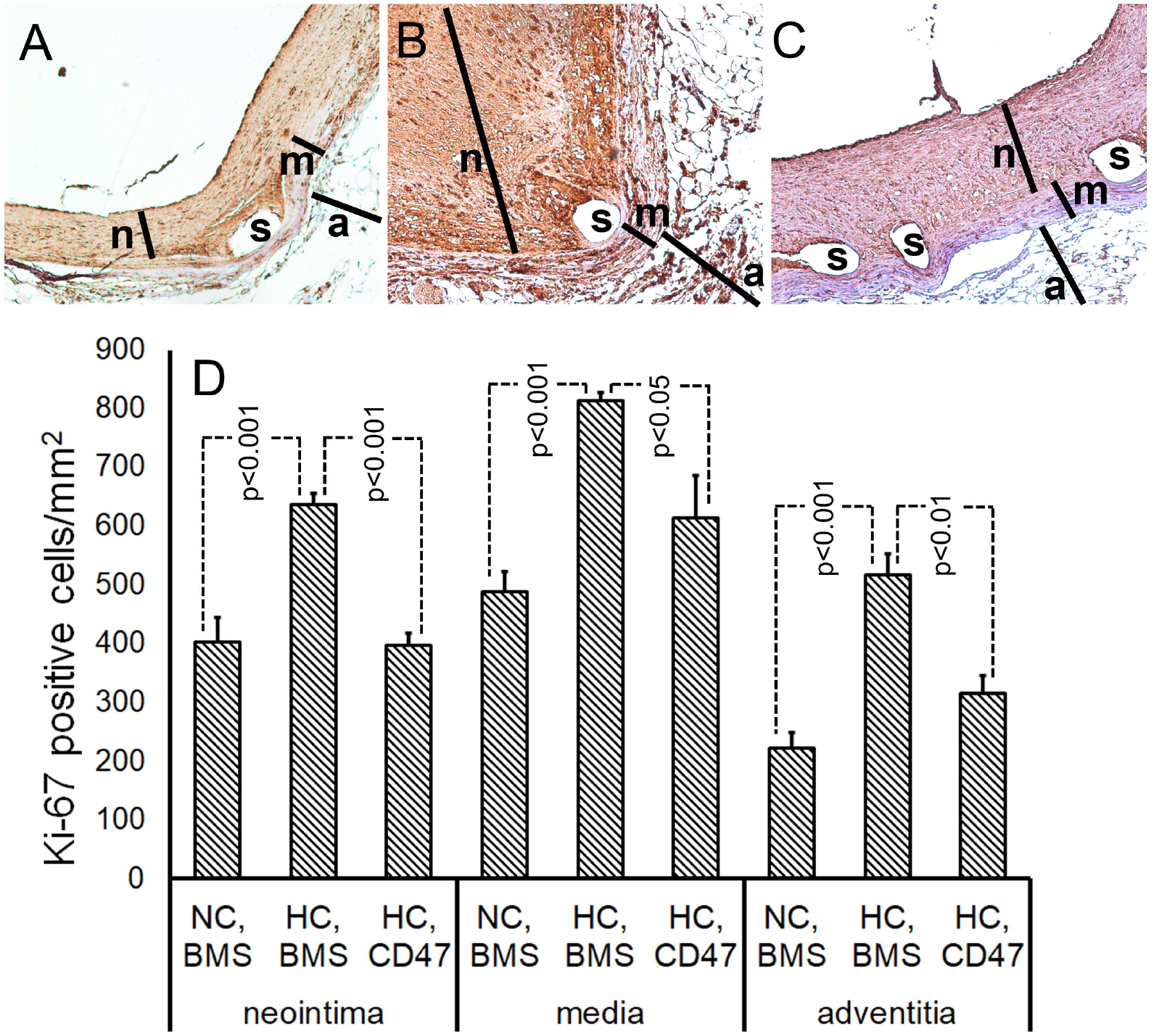
Cell proliferation in the stented arteries. (A-C) Ki67 immunohistochemistry was performed on the sections from the stented iliac arteries of NC (A) and HC (B, C) rabbits treated with BMS (A, B) and pepCD47-modified stents (C). Ki67 labeling was expressed as the number of Ki67-positive cells per mm^2^ of neointimal, medial, and adventitial compartments (D). n – neointima, m – media, a – adventitia, l – lumen, s – strut positions.

## Discussion

### Exacerbation of ISR by hypercholesterolemia is mediated by hypercholesterolemia-induced inflammation

Augmented neointimal response after balloon angioplasty, endarterectomy, and stent implantation was demonstrated in hypercholesterolemic subjects across multiple mammalian species and types of vascular injury^24–27^. Systemic and local arterial inflammation preceding vascular intervention was identified as a pathogenetic link between hypercholesterolemia and restenosis^28–30^. The pro-inflammatory effects of hyperlipidemia are multifaceted. Liver and adipose tissue damage by the excessive dietetic cholesterol triggers the production and secretion of pro-inflammatory cytokines, IL6, IL12, CRP, and TNFα, as well as of chemokines CCL2 and fractalkine (CX3CL1)^8^. In parallel, hypocholesterolemia-induced platelet activation leads to the formation of platelet/leukocyte aggregates that bind avidly to the balloon- or stent-injured arterial wall bridging between the systemic and local vascular inflammation^8^. Inflammation at the site of vascular intervention is further augmented by persisting hypercholesterolemia through the enhanced monocyte accumulation, conversion of macrophages into the foam cell, followed by their death, failed efferocytosis, and activation of NLRP3 inflammasome by the accumulated cholesterol crystals^31^. In accordance with the previous reports^32–34^, we observed increased concentrations of pro-inflammatory cytokines and MDA in the plasma of rabbits fed the HC diet for 8 weeks compared to NC littermates (Fig. 5). Likewise, we confirmed the previously demonstrated exacerbated neointimal growth after stenting in the hypercholesterolemic animals (Fig. 3), and reported on a direct correlation between the total cholesterol levels and ensuing ISR (Fig. 4) that was not demonstrated previously. Plastic embedding techniques typically used for the histological preparation of stented arteries are not routinely compatible with immunohistochemical methods. Therefore very little data about arterial tissue synthesis and secretion of pro-inflammatory mediators are available. *In situ* acid dissolution of stent struts within the explanted arteries^35^ enabled paraffin embedding of “destented” vasculature and subsequent immunohistochemical analysis. Our results (Fig. 6) revealed the peri-strut expression of TNFα, IL1β, and IL6, augmented by persistent hypercholesterolemia (Fig. 6 A, B, and D; E, F and H, and I, J and L, respectively). Likewise, peri-strut macrophage infiltrates detected with the anti-CD68 antibody were significantly more intense in the arteries of the HC compared to the NC animals (Fig. 6 M, N, and P). We have also validated macrophages as the principal source of pro-inflammatory cytokine expression in the stented arteries (Fig. 7). Since the paracrine effect of cytokines secreted by inflammatory leukocytes was firmly established as the main trigger of SMC proliferation in the iatrogenic ally manipulated arteries^23^, we predictably observed increased proliferative activity in all layers of the stented arteries from the HC rabbits compared to their NC counterparts (Fig. 8).

### CD47 functionalization of stents inhibits ISR by blocking early steps of inflammatory pathways

The physiological responses elicited by CD47/SIRPα interaction have become a major topic of interest in cancer research^36^ and transplant biology^37^. Recently, the importance of defective efferocytosis induced by CD47 signaling has been realized as a crucial mechanism of vascular inflammation^38^ and atherosclerotic plaque growth and destabilization^39–41^. Furthermore, a blockage of CD47 signaling either with anti-CD47 antibodies^40^ or a low-molecular-weight inhibitor of the CD47/SIRPs pathway^41^ was shown to be protective against atherosclerosis in the mouse models. Unresolved vascular inflammation is central to the pathogenesis of ISR^42^. To this end, inhibition of CD47/SIRPα signaling was demonstrated to counteract neointimal expansion in wire-injured mouse femoral arteries^43^. Although our previous study^20^ and the present experiments demonstrate the anti-restenotic efficacy of CD47-functionalized stents, our results do not contradict the reports that established the therapeutic effect of CD47 inhibition in comparable animal models of vascular disease. The controversy is explained by the immobilization of the SIRPα-interacting moiety (CD47 peptide^44^) on the stent surface rather than being displayed on the cell membrane. In this presentation, CD47 aborts attachment and spreading of platelets^45^ and cells of myeloid origin^45^, expressing SIRPα. In confirmation of these data, our present studies demonstrated a reduced number of platelets and leukocytes from hypercholesterolemic rabbit blood attached to the pepCD47-modified vs. non-modified stainless steel substrate (Fig. 1). CD47 binds thrombospondin-1^46^, which was implicated in the development of restenosis^47, 48^. Thus, in addition to the anti-inflammatory effects, pepCD47 immobilized on the stent surface may hypothetically sequester thrombospondin-1, limiting its impact on ISR development. Recent studies have highlighted the importance of M1/M2 macrophage equilibrium in the progression of restenosis^49^. Our findings contribute to this growing body of evidence by showing that pepCD47-functionalized surfaces foster the M2 polarization of attached macrophages (Fig. 2).

This observation is in accordance with biased M1 polarization in mouse tumors upon treatment with anti-CD47 antibodies^50^. The underlying mechanism responsible for the CD47-induced M2 polarization is allegedly mediated by the downstream SIRPα signaling through SHP-1 and Akt2^51^.

The ISR severity in hypercholesterolemic rabbits in our study was reduced by 29-52% in the animals treated with pepCD47 stents (Fig. 3), partially offsetting the pro-restenotic consequences of hypercholesterolemia. In agreement with the hypothesis of inflammatory response mitigation with stent-immobilized pepCD47, the macrophage infiltration of peri-strut areas and local arterial expression of pro-inflammatory cytokines in HC animals was reduced to the values observed after stent deployment in the NC rabbit model. The counteraction to the damaging impact of hypercholesterolemia on cytokine expression in the stented vasculature was paralleled by the decrease of the proliferative activity in all layers of the stented arteries (Fig. 8).

In conclusion, the herein reported studies support the view that through systemic and local mechanisms, hypercholesterolemia primes arterial wall to a more vigorous inflammatory response to stenting-triggered injury and, consequently, more robust neointimal formation than observed in normocholesterolemic controls. This untoward outcome is effectively mitigated by pepCD47 immobilized on the surface of stent struts through the inhibition of platelet and leukocyte binding to the stent surface mediated by CD47-SIRPα signaling.

